# Transient demographic approaches can drastically expand the toolbox of coral reef science

**DOI:** 10.1101/2021.04.08.438926

**Authors:** James Cant, Roberto Salguero-Gómez, Maria Beger

## Abstract

Coral communities are threatened by an increasing plethora of abiotic and biotic disturbances. Preventing the ensuing loss of coral coverage and diversity calls for a mechanistic understanding of resilience across coral species and populations that is currently lacking in coral reef science. Assessments into the dynamics of coral populations typically focus on their long-term (*i.e. asymptotic*) characteristics, tacitly assuming stable environments in which populations can attain their long-term characteristics. Instead, we argue that greater focus is needed on investigating the *transient* (*i.e.* short-term) dynamics of coral populations to describe and predict their characteristics and trajectories within unstable environments. Applying transient demographic approaches to the evaluation and forecasting of the responses of coral populations to disturbance holds promise for expediting our capacity to predict and manage the resilience of coral populations, species, and communities.

## Introduction

Coral populations are sensitive to a range of biotic and abiotic pressures (Riegl 2020), many of which will be exacerbated by future climatic change and human exploitation (Wolff et al. 2018; Riegl and Glynn 2020). The future resilience of coral populations subsequently comprises their capacity for enduring, and recovering from, this increased disturbance (Holling 1973). Accordingly, a resilience-orientated focus is becoming commonplace in reef management (Mcleod et al. 2019; McLeod et al. 2021), although this requires the ability for anticipating the responses of coral populations to varying biotic and abiotic regimes (Lam et al. 2020). Appropriately, state-structured demographic models are being increasingly utilised for quantifying the mechanisms supporting or preventing the persistence and revival of disturbed coral communities (e.g. Kayal et al. 2018; Cant et al. 2021). Indeed, understanding how patterns in the survival, size, and reproduction of individual colonies influences population-level characteristics is paramount for effectively managing conservation resources (Riegl et al. 2018; Pisapia et al. 2020). However, the contemporary application of state-structured demographic approaches to ascertain the viability of coral populations does not reflect our need to evaluate their resilience.

Structured population approaches examine how variation at the level of individuals translates to population-level attributes (Caswell 2001). The state of individuals (typically their size, age, or developmental stage [Caswell 2001]) arbitrates their survival and reproduction, which in turn shape the responses of their population to local biotic and abiotic conditions (Benton et al. 2006). Within natural populations, the diversity of individual-level characteristics can be expressed through the state-structured vital rates of survival, progression (growth & development), retrogression (shrinkage [Salguero-Gómez and Casper 2010] & rejuvenation [Salguero-Gómez et al. 2013]), and reproduction (Metcalf and Pavard 2007). Patterns across these vital rates, and their response to changing population structures, subsequently underpin the characteristics of populations and define their capacity for tolerating various environments (Boyce et al. 2006; Ehrlén et al. 2016). Equally, combined with an explicit consideration for the interaction between multiple populations, structured population analyses also facilitate the exploration of community recovery (Kayal et al. 2018) and coexistence (Adler et al. 2010). Evaluating the vital rates and structure of numerous populations, and their association with abiotic regimes, has thus proven instrumental in calculating the extinction risk of highly threatened populations (Mace et al. 2008), and the management, and conservation of natural communities (Morris and Doak 2002; Jongejans et al. 2008).

With many coral communities facing imminent reassembly and population collapse, the demand for an improved understanding of coral population dynamics has never been greater (Edmunds and Riegl 2020). Although limited, the application of demographic theory within coral research is not a novel concept (Edmunds et al. 2014). Veritably, there is a growing appreciation for how the size structure of coral populations influences their dynamics and endurance, with recent appeals for demographic approaches to be concentrated towards gauging the resilience of coral communities (Dietzel et al. 2020; Pisapia et al. 2020; Cant et al. 2021; Lachs et al. 2021). However, prevailing knowledge of coral population dynamics centres around isolated patterns in colony survival, changes in size, or recruitment, and long-term (*asymptotic*) population characteristics. Quantifying the asymptotic characteristics of various populations presents a valuable comparative tool for exploring abiotic tolerances (Beissinger and Westphal 1998). Yet, to achieve their asymptotic dynamics, populations typically require stable conditions, which seldom occur naturally, and are becoming increasingly unlikely given current climate projections (Hastings et al. 2018; Francis et al. 2021). Consequently, a focus towards exploring the asymptotic characteristics of populations limits our appreciation of their dynamics within natural environments (Ezard et al. 2010), and will inhibit our capacity to accurately predict the future condition of global coral assemblages.

Cyclic abiotic and biotic disturbances (e.g. seasonality in temperature or resource availability) ensure that natural environments rarely exist at a stable equilibrium (Williams et al. 2011). The perpetual instability generated by regular disturbances ensures that natural populations rarely attain their asymptotic trajectories (Hastings 2001; Hastings et al. 2018). Instead, the short-term or *transient* characteristics of populations, which reflect their dynamics within non-equilibrate environments, play a fundamental role in the assembly of biological communities (Hastings 2004). With the transient dynamics of populations defining their performance following disturbance, these characteristics also represent quantifiable measures of population resilience (Capdevila et al. 2020). Globally, coral communities are being exposed to increasingly recurrent disturbance events (Hughes et al. 2018), with various species shifting their distributions poleward or establishing populations within marginal habitats (Beger et al. 2014; Camp et al. 2018; Vergés et al. 2019). Here, we discuss why enhancing our capacity for anticipating the future resilience of coral communities requires an understanding of the transient dynamics in their constituent populations.

### Transient *versus* asymptotic population dynamics

Population dynamics characterise and define the trajectories of populations according to the individual-level allocation of resources towards survival, changes in size (or the state of interest), and reproduction (Box 1). Any population can be structured according to the distribution of its individuals across some state variable, be that age, size, developmental stage, or a combination. The transition of individuals across state classes is, in turn, mediated by selection gradients that emerge from the state-specific patterns of survival and reproduction (Groenendael et al. 1988; van Tienderen 2000). In state-structured demography, patterns across these multiple vital rates can subsequently be condensed into a population projection matrix (***A***), with each matrix element (*a_ij_*) expressing the expected transition of individuals into state class *i* from state class *j* during the time interval *t* to *t+1*, or the per-capita a/sexual contributions of state *j* individuals into state class *i* during that same interval (Caswell 2001). This population matrix can then be used to project the size of a population (*N*, number of individuals) over time, using the change in its state structure (**n**):

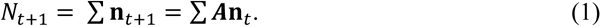

#### Box 1

##### Scaling from individuals to communities

Following individual corals over time facilitates the calculation of vital rate patterns relating to colony survival, size, and reproduction, and how they govern the characteristics of populations and their wider community interactions (Fig. I). As corals mature and grow, they experience changes in their resource demands, with the capacity of corals for continually meeting these changing demands subsequently influencing their ability to maintain their somatic condition and tolerate local biotic and abiotic stressors (Hughes 1984). The repeated survey of individual colonies allows for documenting how the size of individuals regulates their survival, development, and reproductive contribution over time, and how this is shaped by changing environmental conditions. Condensing these temporal observations from across multiple tagged colonies then allows population ecologists to explore how individual-level vital rates underpin the dynamics of populations.

Crucially, state-structured demographic approaches enable the quantification of the relationship between abiotic conditions and population-level characteristics. Such an understanding is necessary if ecologists are to accurately predict the impact of recurrent disturbances and simulate populations under varying climate scenarios. Similarly, by understanding interactions across the dynamics of various co-occurring populations, and their differential responses to changing conditions, ecologists can better understand the mechanisms driving coexistence (Adler et al. 2010), and predict the transition of coral communities amidst changing climatic conditions (Hughes 1996; Kayal et al. 2018; Cant et al. 2021). Indeed, although more time consuming, demographic approaches transcend the correlative techniques previously used for evaluating the viability of coral populations and communities, and for predicting their resilience to future climatic stressors (Edmunds et al. 2014; Edmunds and Riegl 2020).

**Figure I.**
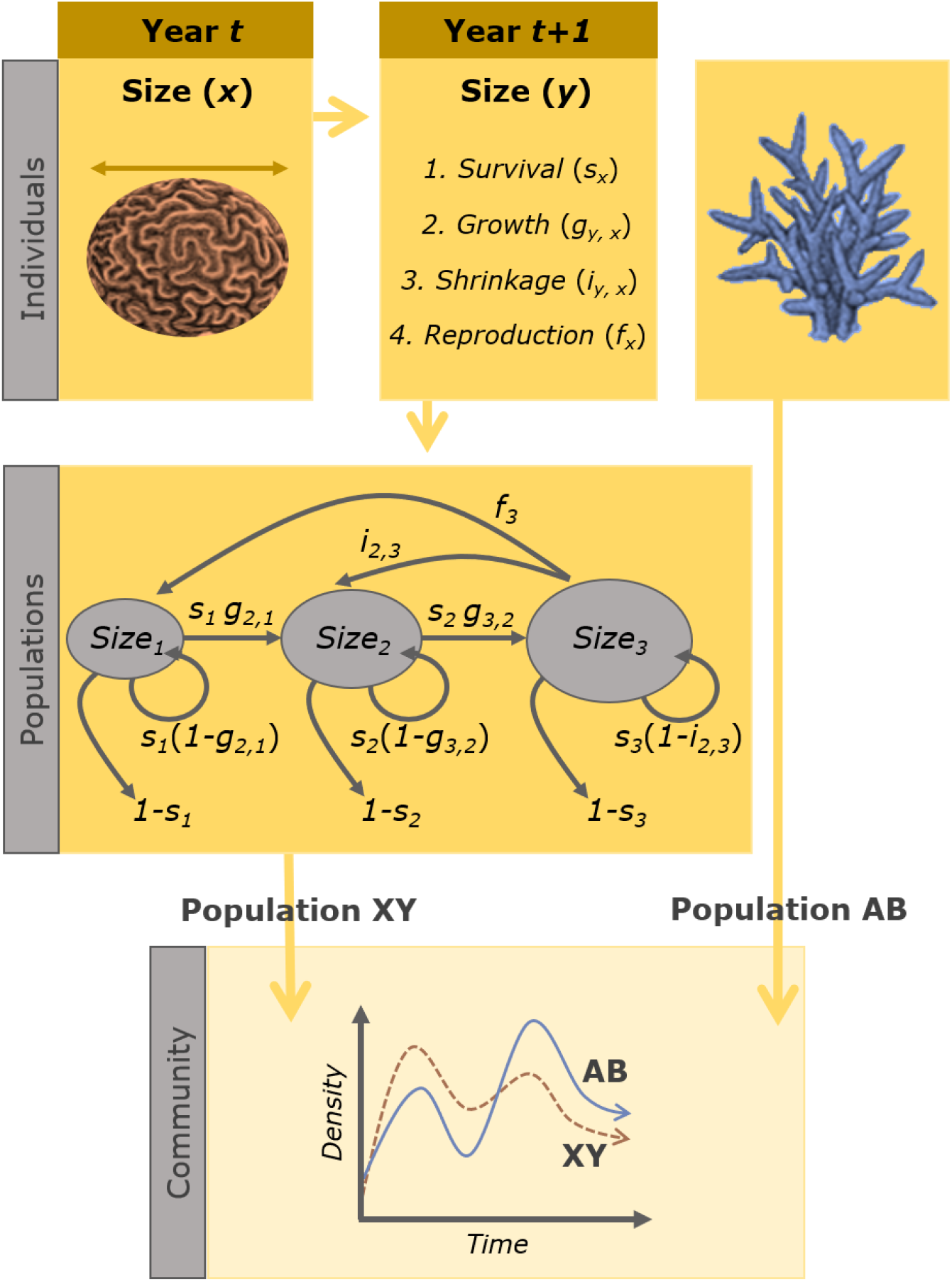
Records of survival, size, and reproductive output, collected from tagged coral colonies over time, can be used to enhance predictions of population trajectories and community reassembly under future climate scenarios. Here, subscript notation is used to reflect how the changing state of individuals corresponds with changes in individual-level characteristics. In this schematic the state of individuals refers to colony size and thus the growth (*g*) of individuals between size *x* and *y* is expressed as *g*_*yx*_.

Within a stable environment, the state structure of a population is expected to converge towards an equilibrium state (Caswell 2001). At equilibrium, the size of a population is then expected to change at a constant rate over time, termed its asymptotic population growth rate (*λ*, Caswell 2001). The value of *λ*, for any state-structured population, is equal to the dominant eigenvalue of its corresponding population matrix ***A***. Estimates of *λ* represent the proportional change in the size of populations over time, and exist on a non-negative scale reflecting population growth (*λ* > 1), decline (*λ* < 1), or stasis (*λ* = 1).

Crucially, though, with populations rarely exposed to stable conditions or equilibria, deterministic estimates of *λ* are unlikely to represent a true reflection of a population’s trajectory (Boyce 1992; Beissinger and Westphal 1998). To achieve their asymptotic growth rate populations require a stable population structure, at which their vital rate patterns will maintain a consistent distribution of individuals across state classes over time (Caswell 2001). Yet, for their stable structure to manifest, populations typically require time in a constant environment to allow the state distribution of individuals to convergence towards equilibrium, without displacement by local disturbances (Fig 1; Bierzychudek 1999). Additionally, the vital rate patterns used in constructing population projection matrices are themselves not consistent, and contingent on the environmental conditions in which they are observed (Benton and Grant 1996; Boyce et al. 2006). Equally, as populations change in size their dynamics are subject to density–dependant processes that regulate their trajectories within shared environments (Engen et al. 1998; Lande et al. 2003). Thus the dynamics of populations can be expected to vary over time as environmental conditions change (Tuljapurkar and Orzack 1980; Tuljapurkar 1989).

**Figure 1.**
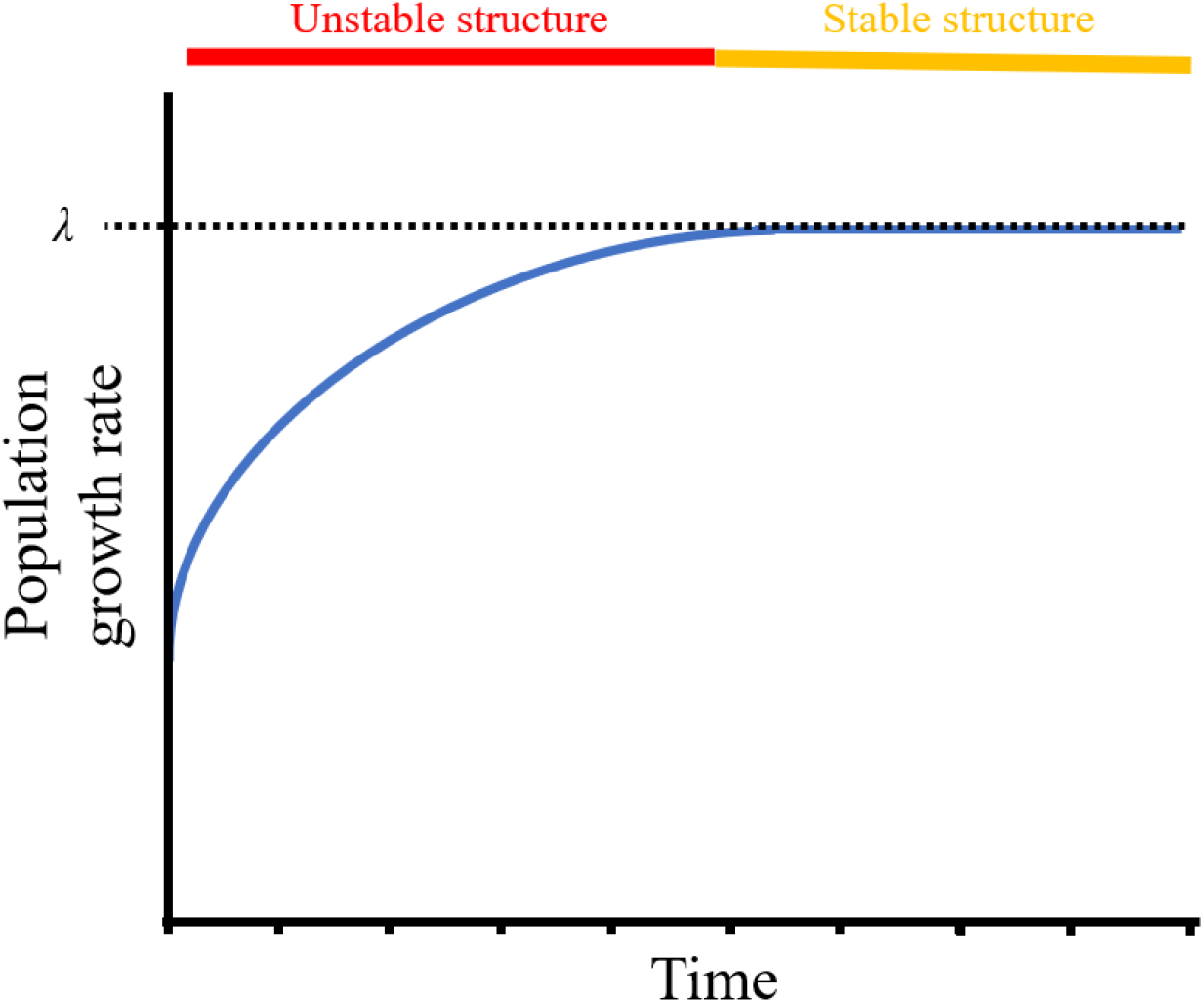
At stationary equilibrium, populations are expected to change size at a constant rate, termed their asymptotic growth rate (λ). However, to display asymptotic characteristics, populations need a stable state structure and so require time at equilibrium to enable the development of this stable structure. In the absence of a stable state structure, populations will display varying growth rates as they converge towards asymptotic expectations, even within equilibrate environments. Note that, although in this schematic the populations growth rate is increasing towards its asymptotic trajectory, estimates of λ can also reflect asymptotic decline.

Recurrent perturbations prevent populations from acquiring, or preserving, a stable state structure (Bierzychudek 1999; Williams et al. 2011). Instead, environments persist within a non-equilibrate condition maintaining populations within a transient state during which their trajectories can radically differ from asymptotic expectations (Stott et al. 2010, 2011; Hastings et al. 2018). Accordingly, anticipating the future condition and endurance of populations requires an awareness for their transient characteristics (Box 2; Ezard et al. 2010). Moreover, with the transient characteristics of populations reflecting their response to disturbance and instability, they represent quantifiable measures of population resilience (Capdevila et al. 2020). Transient shifts in the state structure of populations can elevate (*amplification*) or diminish (*attenuation*) their growth rates relative to asymptotic projections (Townley et al. 2007). Equally, for populations existing in a transient state, there is a natural tendency for their state structure to converge towards a stable structure (Caswell 2001). Intuitively, with population resilience comprised of the attributes of *resistance* (the ability to withstand disturbances), *compensation* (the ability to benefit from disturbances) and *recovery* (the speed and ability to reattain stability), the transient characteristics of populations resemble these features of resilience (Hodgson et al. 2015; Capdevila et al. 2020).

#### Box 2

##### Measuring transient population characteristics.

Within natural environments, abiotic and biotic perturbations generate constant shifts within the state structure of populations ensuring they persist within a transient state, rather than converging towards equilibrium (Hastings et al. 2018). Within this transient state, the growth rate of populations can differ dramatically from their asymptotic trajectories, with populations undergoing *amplification* (increases in growth rate) or *attenuation* (declines in growth rate), which can subsequently influence the viability of a population in the longer-term (Fig. II). Accordingly, it is necessary to explore the transient characteristics of populations to gain insights into their responses to disturbance, and therefore better understand their resilience (Ezard et al. 2010; Capdevila et al. 2020). Evaluating the transient dynamics of populations is made possible through the calculation of transient indices from population projection matrices (Stott et al. 2011). These measures include population *reactivity* 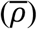 and *first-timestep attenuation* (*<u>ρ</u>*) which describe the increase or decline in a populations growth rate within one time interval of a perturbation, relative to *λ* (Stott et al. 2011). Equally, the measures of *maximal amplification* 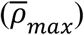 and *maximal attenuation* (*<u>ρ</u>*_*max*_) can be used to evaluate the maximum amplification or attenuation expected in the growth rate of populations, relative to *λ* (Townley et al. 2007; Townley and Hodgson 2008).

The *damping ratio* (*ρ*) of a population defines the rate at which at population will converge back to a stable equilibrium following a perturbation to its stable structure (Caswell 2001). Finally, *upper* & *lower population inertia* (*Inertia*_*upper*_ & *Inertia*_*lower*_ respectively) can be used to explore the extent to which the transient characteristics of populations influence their long-term trajectories, also known as a populations *transient envelope* (*TE*) (Koons et al. 2005; Stott et al. 2011).

**Figure II.**
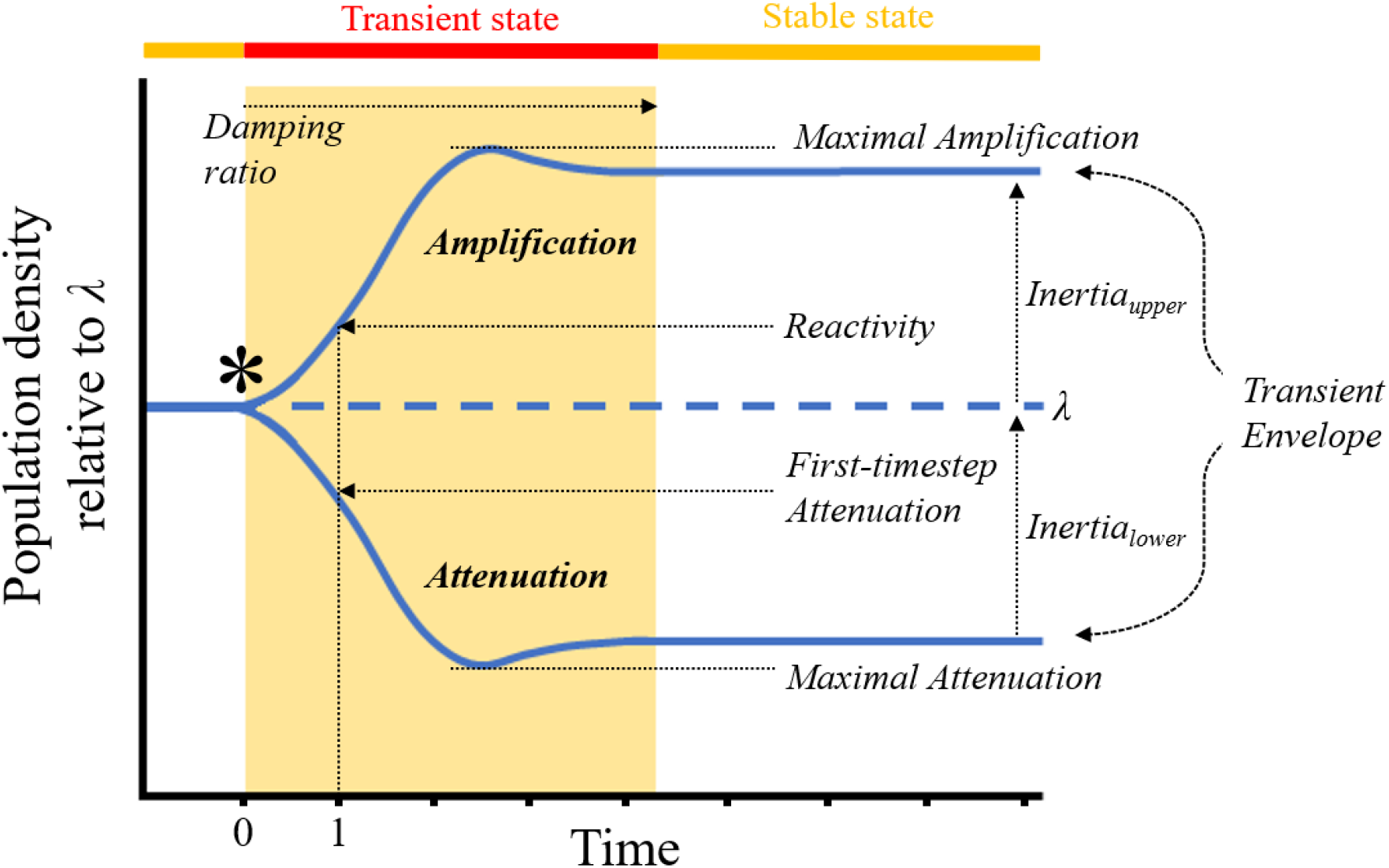
Various measures exist for quantifying the transient characteristics of populations, and how the short-term responses of populations following a disturbance (*) will modify their trajectories relative to asymptotic expectations.

Evaluating the transient characteristics of populations, and therefore their demographic resilience, is possible through the calculation of transient measures readily accessible from population projection matrices (Box 2; see Stott et al. [2011] for a detailed review). Briefly, measures of *reactivity* 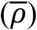 and *first-timestep attenuation* (*<u>ρ</u>*) reflect the initial behaviour in a population’s growth rate following a perturbation, with the *damping ratio* (*ρ*) describing the rate at which a population is expected to reacquire a stable state structure (Caswell 2001; Stott et al. 2011). Alternatively, estimates of *maximal amplification* 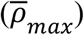, *maximal attenuation* (*<u>ρ</u>*_*max*_), and *population inertia* (*Inertia*_*x*_) illustrate the magnitude by which the growth rate of a population can be expected to fluctuate, and the influence of this short-term variation on the population’s long-term trajectory (Koons et al. 2005; Townley et al. 2007; Stott et al. 2011). Furthermore, with population projection matrices providing a link between each transient characteristic and a population’s vital rates, sensitivity analyses can be applied to evaluate the mechanistic drivers underlying the short-term characteristics of populations (Stott 2016; Caswell 2019). Overall, transient demographic approaches present an effective tool for explicitly evaluating the resilience of natural populations (Capdevila et al. 2020), and have proven instrumental in our understanding of the establishment of populations within new and variable environments (Iles et al. 2016; McDonald et al. 2016, 2017; Jelbert et al. 2019).

### Coral population dynamics: Current knowledge

To examine the advancement of state-structured population approaches within coral research, we conducted a review of the relevant ecological literature. Specifically, we searched for the term coral* AND demograph* across the titles, abstracts, and keywords of research literature within the SCOPUS database. We applied our search across all indexed peer-reviewed journals published between 1960 and 2020, inclusive. This search initially returned 781 studies, from which we retained all studies that explicitly measured or quantified state-structured patterns in the survival, size transitions (including shrinkage via partial mortality or fission), and/or reproduction (e.g. fecundity) of coral colonies. We also retained studies measuring larval settlement rates, as the recruitment of early life stage individuals is a key demographic aspect within coral communities (Adjeroud et al. 2017). Following this refinement, we retained a total of 145 studies (~19% of our original search), which we then categorised according to whether they involved the presentation of (i) asymptotic dynamics/simulations, (ii) transient characteristics, or (iii) an assessment of vital rate patterns only.

Following the seminal work of Hughes and Jackson (1980), the application of state-structured demographic assessments has increased within coral research, with a particularly rapid growth in popularity since 2007 (Fig. 2). However, much of this research (97%) has focused on either exploring the asymptotic dynamics of coral populations (34%; 50 studies) or solely evaluating patterns across one or more of the vital rates of survival, changes in size, or reproduction (63%; 91 studies). Presently, the majority of knowledge regarding the dynamics of coral populations centres around their long-term trajectories. Thus, our current understanding regarding the dynamics of coral populations assumes that these populations will experience the stable environments necessary for achieving optimum population structures.

**Figure 2.**
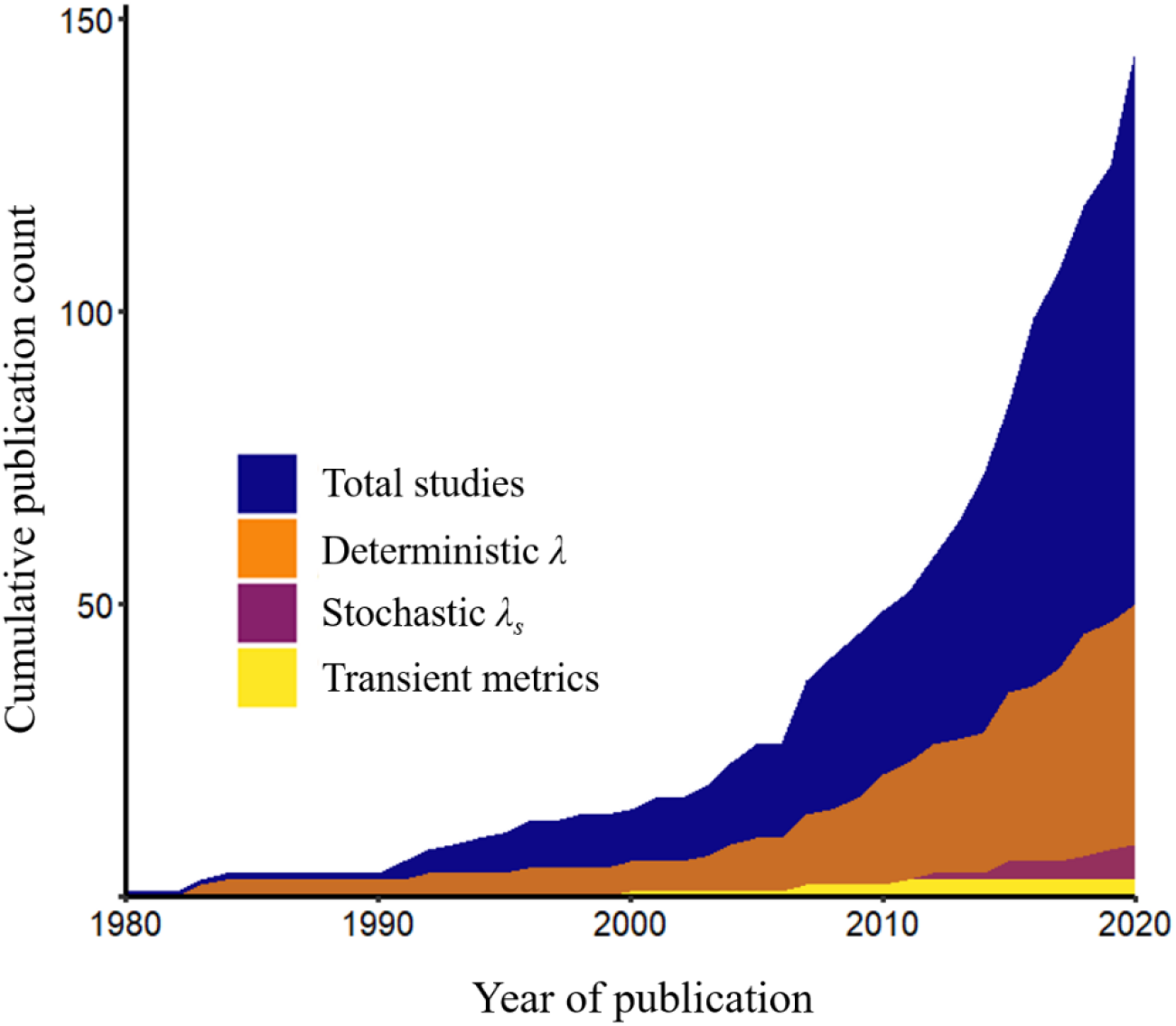
The cumulative number of studies that explicitly explore the state-structured demographics of coral populations as identified by our search for the term coral* AND demograph* across the peer-reviewed literature within the SCOPUS database. The colour scale differentiates between studies that focus on only investigating patterns in colony survival, growth, and/or reproduction, and those that utilised estimates of either deterministic population growth (*λ*), stochastic asymptotic characteristics (*λ*_*s*_), or transient population characteristics, to evaluate the dynamics of coral populations.

A limited amount of previous research has considered the unpredictable nature of natural environments when evaluating the future trajectories of coral populations. Our search identified nine studies that either included estimates of stochastic population growth (*λ*_*s*_; Lewontin and Cohen 1969) or explicitly parameterised the density-dependant regulation of vital rates within their population models (Fig. 2; see Linares et al. 2007; Linares and Doak 2010; Hernandez-Pacheco et al. 2011; Vardi et al. 2012; Bramanti et al. 2015; Mercado-Molina et al. 2015; Kayal et al. 2018; Montero-Serra et al. 2019). Estimates of *λ*_*s*_ describe the average periodic change in the size of a population over a series of successive time intervals (Engen and Saether 1998). As such, this metric of population performance provides a more accurate consideration of vital rate and environmental variation when evaluating future population trajectories, compared to deterministic *λ*. However, with estimates of *λ*_*s*_ often derived using a series of population sizes estimated over numerous time intervals or with initial inconsistent estimates of population size omitted, *λ*_*s*_ is still fundamentally a long-term measure of population viability (Ellner and Rees 2007). Consequently, this measure still neglects the short-term characteristics of populations, and thus our capacity to fully anticipate the future dynamics of coral populations remains restricted.

Whilst state-structured demographic approaches are being adopted within coral research, our review of the literature demonstrates how prominently the assumption of stable equilibria dominates our understanding of coral population dynamics. Subsequently, we lack the understanding necessary for accurately projecting the future reassembly and persistence of coral communities under increasingly recurrent disturbance regimes. Only three of the 145 studies in our literature search calculated measures of the transient characteristics of coral populations (Fig. 2; see Hughes and Tanner 2000; Linares et al. 2007; Bruno et al. 2011). Alongside asymptotic population characteristics, these studies estimated population damping ratios to outline the rate at which their focal populations would converge back to a stable state structure following a perturbation. Yet, the damping ratio represents a dimensionless, mathematical property of population projection matrices, and alone is of limited use to population managers seeking to predict the short term responses of populations to disturbance (Stott et al. 2011). Forecasting the resilience of coral populations is essential for the future conservation of various reef communities (Roche et al. 2018; Lam et al. 2020). We have illustrated here, however, that current approaches used in coral population ecology to evaluate population growth rates do not constitute measures of population resilience. Our understanding of coral population dynamics needs to be further expanded in order to meet our need for quantifying the resilience, and responses, of coral communities to increasingly frequent disturbances.

### Transient dynamics and the resilience of coral communities

Exploring the transient characteristics of coral populations will enhance our understanding regarding the future diversity, condition, and resilience, of coral communities. The frequency of widespread thermal stress events is increasing across global coral communities, and soon expected to exceed the recovery time required by most coral species (Hughes et al. 2018; Sully et al. 2019). Presently, considerable value is placed on classifying coral species according to their morphological and functional traits, with the subsequent categorisation of coral taxa offering proxies for the vulnerability of various coral communities to said recurrent stress events (Darling et al. 2012, 2019). Quantifying the relative affinities of different coral populations for demographic recovery presents the opportunity for justifying these assumptions; helping to distinguish the species and populations most vulnerable and or resilient to projected recurrent disturbance regimes.

Intuitively, the amplification and attenuation characteristics of populations simultaneously reflect their stability following a disturbance. However, recognising the differential attributes of compensation and resistance in different populations is important for managing the future condition of biological communities (Capdevila et al. 2020). Assessments of coral communities following repeated disturbances have demonstrated divergent shifts in species compositions (Adjeroud et al. 2009; Pratchett et al. 2011, 2020). With knowledge of the capacity for various coral populations to undergo demographic amplification or for avoiding demographic attenuation, we can begin to forecast and anticipate the reassembly of coral communities, thereby improving the effectiveness of adaptive reef management. Alternatively, a disturbance does not necessarily imply a negative impact, merely a change to a system (Pickett and White 1985). Disturbances to the structure of populations can be brought about by the cessation of a continuous pressure, changes in resource availability, or establishment within a novel environment. The transient characteristics of populations offer insights into their invasive potential (Iles et al. 2016), with the ability of natural populations for undergoing demographic compensation underpinning their capacity for exploiting changes within their local environments (Jelbert et al. 2019). Accordingly, comprehending and quantifying demographic compensation within coral populations will aid predicting the winners and losers in reef communities targeted by conservation initiatives, restoration strategies, or climate induced range shifts. Assessing coral reef community resilience within such a framework requires an increased focus on the temporal nature of resilience, a view that would transcend current approaches of conducting single reef assessments to determine resilience (Maynard et al. 2015; McLeod et al. 2021).

Complexities in the modelling approaches used to explore the dynamics of natural populations have resulted in these techniques remaining largely overlooked within coral research (Edmunds et al. 2014). Indeed, parametrising the demographic models needed to quantify population characteristics requires considerable amounts of data (Ellner et al. 2002). Although, despite the data demanding nature of state-structured demographic models, the collection of the necessary data regarding the survival, transitions in size, fragmentation, and recruitment, of individual coral colonies is possible alongside current reef monitoring efforts (Edmunds and Riegl 2020). Furthermore, new techniques for analysing the demographic characteristics of populations structured by a continuous state variable like size (Integral Projection Models [IPMs; Easterling et al. 2000]) have reduced the data demands of demographic assessments (Ramula et al. 2009). As a result, IPM frameworks are growing in popularity throughout coral research (e.g. Bruno et al. 2011; Madin et al. 2012; Zychaluk et al. 2012; Elahi et al. 2016; Kayal et al. 2018; Precoda et al. 2018; Scavo Lord et al. 2020; Cant et al. 2021; Carlot et al. 2021). Finally, studying the dynamics, particularly the transient characteristics, of coral populations is made increasingly accessible by the development of demographic analysis packages such as *popdemo* (Stott et al. 2012).

It is not our intention to dismiss previous efforts to evaluate the dynamics of coral populations using asymptotic characteristics. Changes in individual vital rates can of course have critical repercussions on the dynamics of populations (e.g. Bellier et al. 2018), and asymptotic characteristics provide valuable insights for the management of threatened populations (e.g. Johnson et al. 2010). However, as we have illustrated here, the asymptotic characteristics of populations do not represent indicators of population resilience. Here, we advocate for a greater commitment towards investigating the transient dynamics of coral populations that more accurately reflect their dynamics within natural environments (Ezard et al. 2010; Stott et al. 2011; Capdevila et al. 2020). With changing climatic regimes and increased anthropogenic interference exposing global reef ecosystems to increasingly frequent disturbances (Hoegh-Guldberg et al. 2017), evaluating the transient dynamics of coral populations, and their mechanistic drivers, will provide a new effective lens on coral reef resilience.

## Acknowledgements

The authors would like to thank D. Ceccarelli for her comments on early drafts of this manuscript. This work was supported by a NERC DTP scholarship to JC, and a NERC Independent Research Fellowship (NE/M018458/1) to RS-G.

